# The FLAME-accelerated Signalling Tool (FaST): A tool for facile parallelisation of flexible agent-based models of cell signalling

**DOI:** 10.1101/595645

**Authors:** Gavin Fullstone, Cristiano Guttà, Amatus Beyer, Markus Rehm

## Abstract

Agent-based modelling is particularly adept at modelling complex features of cell signalling pathways, where heterogeneity, stochastic and spatial effects are important, thus increasing our understanding of decision processes in biology in such scenarios. However, agent-based modelling often is computationally prohibitive to implement. Parallel computing, either on central processing units (CPUs) or graphical processing units (GPUs), can provide a means to improve computational feasibility of agent-based applications but generally requires specialist coding knowledge and extensive optimisation. In this paper, we address these challenges through the development and implementation of the FLAME-accelerated signalling tool (FaST), a software that permits easy creation and parallelisation of agent-based models of cell signalling, on CPUs or GPUs. FaST incorporates validated new agent-based methods, for accurate modelling of reaction kinetics and, as proof of concept, successfully converted an ordinary differential equation (ODE) model of apoptosis execution into an agent-based model. We finally parallelised this model through FaST on CPUs and GPUs resulting in an increase in performance of 5.8× (16 CPUs) and 53.9× respectively. The FaST takes advantage of the communicating X-machine approach used by FLAME and FLAME GPU to allow easy alteration or addition of functionality to parallel applications, but still includes inherent parallelisation optimisation. The FaST, therefore, represents a new and innovative tool to easily create and parallelise bespoke, robust, agent-based models of cell signalling.

## Introduction

Cellular signalling is essential in translating extrinsic and/or intrinsic chemical and physical stimuli into diverse cell responses such as proliferation, cell migration or cell death. The duration of stimuli, concentration of stimuli, concentration of signalling components, the reaction kinetics in signalling pathways and subcellular localisation of components can drastically affect downstream outcomes of cell signalling pathways. Moreover, cell-signalling pathways, typically, are highly complex with redundancy, cross-talk between different signals and numerous levels of regulation complicating our understanding of how cellular decisions are made. Systems biology approaches have increasingly been used to better understand and predict outcomes from cell signalling processes [1,2]. The most commonly used approach is ordinary differential equation (ODE) modelling that uses a series of differential equations to define how the concentrations of reactants change over time. This has been used effectively to describe a number of cell signalling pathways, including the NFκB pathway [3,4], the intrinsic apoptosis pathway [5–7] and the cell cycle [8]. However, biological systems are characterised by, complex structural organisation, a great level of heterogeneity and physical phenomena, such as molecular crowding, that are not adequately included in ODE models. Furthermore, stochastic effects in biological settings can have profound knock-on effects on cell signalling outcomes [9,10]. Whilst, efforts have been made to better consider all these effects in ODE models [11,12] and by using other modelling techniques such as partial different equation (PDE) models [13–15] or delay differential equation (DDE) models [16], improved methods are required for deeper understanding of complex nanoscale and system level events in cell signalling.

Agent-based modelling (ABM) is a type of bottom up systems modelling that has recently gained popularity in the study of cell signalling pathways and other biological processes [17]. ABM of cell signalling models behaviour of individual molecules and their interactions at the nanoscale. ABM is a powerful tool for modelling cell signalling as complex geometry is easily included and behaviour is naturally stochastic. However, ABM is computationally prohibitive, as the actions and interactions of potentially millions of individual signalling molecules over considerable periods of times must be considered. Furthermore, for ABM to be truly reflective of the modelled system it should be able to robustly model reaction kinetics. A number of ABM methods have been applied to cellular signalling previously, also giving rise to formal simulators such as Smoldyn, ChemCell and MCell [18–23]. These simulators offer highly robust and user-intuitive ABM of cell signalling pathways but still contain limits in scale-up of simulations, as well as, flexibility in their manipulation beyond the inbuilt functionality.

Parallel computing, the distribution of computational work across multiple central processor units (CPUs) or on graphical processing units (GPUs), offers the possibility to improve scale up of ABM simulations. FLAME (Flexible Large-scale Agent-based Modelling Environment) and FLAME GPU are generalised ABM platforms that are used to create ABM applications that can be easily parallelised on CPUs and GPUs respectively. FLAME and FLAME GPU use a communicating X-machine approach to parallelisation, where the user declares discrete functions with input and output communication messages in an XMML file and then the functions are declared in C. This allows FLAME and FLAME GPU to build the discrete functions into a parallel model therefore removing the necessity of user knowledge of message passing interface (MPI) or CUDA coding respectively [24,25]. Furthermore, they contain intrinsic parallelisation optimisation, even when including new functionality (for a short summary of FLAME’s approach to parallelisation, see supplementary S1, for a full technical report of FLAME’s and FLAME GPU’s approaches to parallelisation, see the reports in [24,25]).

In this paper we establish and validate new methods for accurate ABM of cell signalling. We implement these methods into the FaST (FLAME-accelerated Signalling Tool), which creates ABM models from reaction networks that can be easily customised and parallelised on CPUs or GPUs using FLAME and FLAME GPU. We then demonstrate that this tool can convert a previously established ODE model of apoptosis execution into an ABM simulation that reliably reproduces the kinetics of the original ODE model. Moreover, the performance of this simulation could be vastly improved by CPU parallelisation and GPU-acceleration.

## Results

### Simulation of the random walk

In order to establish methods for ABM of cell signalling, we started by focusing on the movement of individual molecules within suspension. Particles in suspension undergo Brownian motion, a random walk caused by collisions with molecules of the solvent [16]. This can be simulated by implementing the polar form of the Box-Muller transformation of uniformly distributed random numbers into normally distributed random numbers [26,27]. These are then scaled to fit the Gaussian distribution for the change in *x*, *y* or *z* (*Δx*, *Δy*, *Δz*):

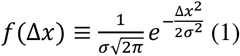

where *σ* is calculated from the translational diffusion coefficient *D*_*T*_ in m^2^⋅s^−1^ and the time step *Δt* in s:

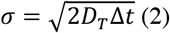

as demonstrated previously [18,28]. Three particles, with diffusion coefficients of 1 μm^2^⋅s^−1^, 5 μm^2^⋅s^−1^ and 10 μm^2^⋅s^−1^, reflective of diffusion coefficients of proteins in biological membranes and under molecular crowding [29–31], were simulated for 5 minutes and the 3D traces are shown in Fig 1a. The implementation of Brownian motion was assessed using the mean squared displacement (MSD):

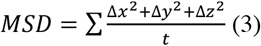

which is related to the diffusion coefficient, such that:

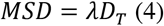

where *λ* is a constant of dimensionality equal to 2, 4 or 6 for 1D, 2D and 3D respectively. The observed MSD was compared to the expected MSD calculated with Equation 4 in Fig 1b. The observed MSD shows excellent agreement with the expected MSD over the 5 minute simulation with observed diffusion coefficients calculated from Equation 4 of 1.00 μm^2^⋅s^−1^, 4.99 μm^2^⋅s^−1^ and 9.99 μm^2^⋅s^−1^.

**Fig 1.**
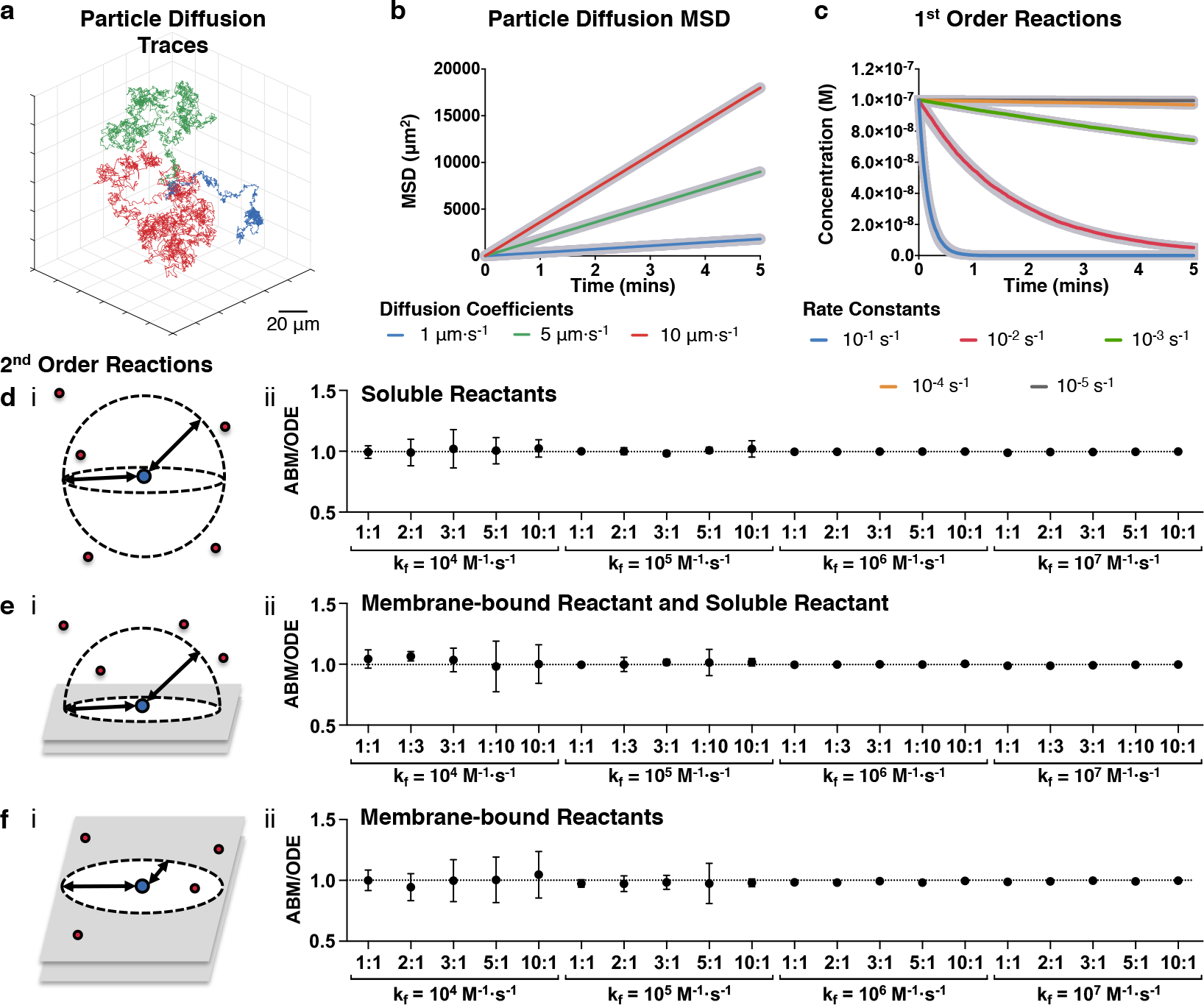
Agent-based models are able to reproduce mass action kinetics. Particles with diffusion coefficients of 1 μm^2^⋅s^−1^ (blue), 5 μm^2^⋅s^−1^ (green) and 10 μm^2^⋅s^−1^ (red) were simulated **(a)** and the observed MSD was compared to the expected MSD (thick grey lines) **(b)**. First order reactions with *k* values of 10^−1^ s^−1^, 10^−2^ s^−1^, 10^−3^ s^−1^, 10^−4^ s^−1^ and 10^−5^ s^−1^ were simulated for 5 minutes with a time step of 0.05 s by ABM (thin lines) and compared to equivalent ODE models (thick grey lines) **(c)**. The second order reactions of two soluble molecules **(d)**, a soluble molecule with a membrane-bound molecule **(e)** and two membrane-bound molecules **(f)** were simulated with ABM and the ABM:ODE Score between the ABM simulations and equivalent ODE models were plotted for *k*_*f*_ alues of 104 M^−1^⋅s^−1^, 10^5^ M^−1^⋅s^−1^, 10^6^ M^−1^⋅s^−1^ and 10^7^ M^−1^⋅s^−1^ and different indicated concentration ratios of A to B ([A]:[B]). Dashed lines indicate the perfect agreement between ABM and ODE of 1. All simulations were for 5 minutes, the time step *Δt* for particle diffusion in all simulations was 0.0001 s and for reactions was 0.05 s. The diffusion coefficients used in **d-f** were 30 μm^2^⋅s^−1^ for soluble molecules and 0.3 μm^2^⋅s^−1^ for membrane-bound molecules.

### Simulation of first order reactions

First order reactions, such as degradation, dissociation and catalysis, form integral parts of cell signalling pathways. Therefore, we next set out to establish and validate methods to simulate first order reactions by ABM. First order reactions of the forms:

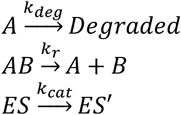

can be simulated by calculating a probability *P* that a single molecule will react within a single discrete time step of time *Δt*, equal to:

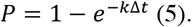

A simulated molecule will react if a randomly generated number is less than the probability calculated by Equation 5 from its own *k* value, given in s^−1^. The degradation of 100 nM of molecule *A*, with *k*_*r*_ values of 10^−1^ s^−1^, 10^−2^ s^−1^, 10^−3^ s^−1^, 10^−4^ s^−1^ and 10^−5^ s^−1^, was simulated by this method and compared to the expected kinetics from the ODE rate equation:

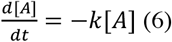

in Fig 1c. Fig 1c shows excellent agreement with the expected kinetics generated from the ODE reaction with an almost exact overlay, thus demonstrating that ABM can effectively model first order reactions.

### Simulation of second order reactions

Many important biological reactions can be described by second order reaction kinetics where two molecules react together. Therefore, we next looked to establish and validate ABM methodology describing second order reactions of the form:

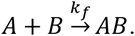

### Second order reactions of two soluble reactants

Pogson and colleagues previously described a method for ABM of second order reactions that is valid when both reactants are freely soluble [22]. A molecule of *A* reacts with a molecule of *B* if the molecule of *B* ends the iteration within an interaction volume *V*_*i*_ around *A*, calculated from the *k*_*f*_ in M^-1^⋅s^-1^ and the time step *Δt*. Assuming *V*_*i*_ is distributed as a sphere around the centre of mass of *A* then the two molecules react if they end an iteration separated by less than an interaction distance *d*_*i*_:

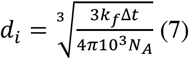

where *N*_*A*_ is Avogadro’s constant (Fig 1d.i). Full derivation of the *V*_*i*_ and *d*_*i*_ is well described and illustrated in the publication of Pogson and co-workers.

### Second order reactions of a membrane-bound reactant and a soluble reactant

Whilst Pogson and colleagues consider reactions involving two soluble reactants, they do not explicitly address the interaction of a soluble reactant with a membrane-bound reactant. However, these types of reactions are often an integral part of many cell-signalling pathways, for example, in receptor-ligand binding. Therefore, we next set out to extend these methods to include this type of second order reaction. In reactions involving a membrane-bound and a soluble reactant, in most cases, the soluble reactant is unable to freely cross the membrane. Consequently, by assuming a sufficiently small *V*_*i*_ limits the impact of membrane curvature, it can be assumed that *V*_*i*_ is distributed as a hemisphere around the receptor (Fig 1e.i). The interaction distance *d*_*i*_ for *R* to react with *S* is therefore equal to:

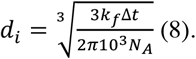

### Second order reactions of two membrane-bound reactants

Membrane-bound molecules also can participate in second order reactions, for example, in receptor clustering and dimerisation. Therefore, we next established ABM methodology for the second order reaction of two membrane-bound reactants. When considering the reaction of two membrane-bound molecules the general principles are the same as for soluble interactions except that as reactions take place on a planar membrane, molecules have an interaction area *A*_*i*_ rather than an interaction volume (Fig 1f.i). In order to hold true for receptor-receptor interactions, receptor levels should be measured in density with units m^-2^ and *k*_*f*_in m^2^⋅s^−1^ giving an interaction distance *d*_*i*_ of:

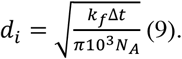

However, in many cases concentration is still measured in M and the *k*_*f*_ determined using soluble forms of the membrane protein in the units of M^−1^⋅s^−1^. In these situations a modified value *k*_*f*_, called *k_f_’*, can be calculated that is scaled to the surface area to volume ratio. In internal cellular reactions this is the cell membrane surface area *A*_*c*_ and the cell cytosolic volume *V*_*c*_ so that:

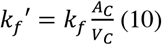

The substitution of Equation 10 into Equation 9 gives the interaction distance when *k*_*f*_ is in the units of M^−1^•s^−1^ as:

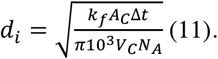

### Agent-based modelling can reproduce mass action kinetics for second order reactions

In order to validate that these methods for ABM of second-order reactions are capable of reproducing mass action kinetics, we conducted a series of simulations by ABM for all three methods and compared these to equivalent ODE models. The rate equations used were:

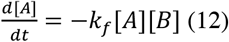

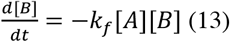

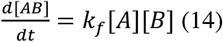

with different *k*_*f*_ values (10^4^ M^−1^⋅s^−1^, 10^5^ M^−1^⋅s^−1^, 10^6^ M^−1^⋅s^−1^, 10^7^ M^−1^⋅s^−1^) and different ratios of reactants. The progress of each reaction was plotted against time for soluble-soluble (supplementary S2 Fig), membrane-soluble (supplementary S3 Fig) and membrane-membrane reactions (supplementary S4 Fig). These figures show a good overlay of the ABM over the ODE curves for each of the second order reaction methods described across a range of different conditions. We then went further in numerically assessing the accuracy of each individual ABM simulation by taking the ratio of the concentration of the product, *AB*, in ABM simulations ([*AB(t)*]_*ABM*_) against the concentration in ODE simulations ([*AB(t)*]_*ODE*_) at individual time points ([*AB(t)*]_*ABM*_:[*AB(t)*]_*ODE*_). We did this every 0.05 s for 5 minutes, or until reaction completion, and then calculated the average ratio (ABM:ODE Score) as a score with an idealised value of 1 representing perfect agreement. Each reaction was repeated in three independent ABM simulations and the calculated ABM:ODE Scores were plotted in Figs 1d.ii, 1e.ii, and 1f.ii for two soluble reactants, a membrane-bound to a soluble reactant and two membrane-bound reactants reactions respectively.

The data in Figs 1d-f show good agreement between the ABM and ODE with all mean values centred on, or proximal, to the perfect agreement ratio of 1. When the reaction is slower, due to the low *k*_*f*_ value and low levels of reactants, the amount of noise increases due to natural stochastic effects having greater weight relative to the mean. However, the mean values still demonstrate excellent agreement with the ODE even under these conditions. It may be expected that when the binding interaction distance and concentration of reactants is high that these methods will undervalue the reaction kinetics because of the increasing probability of multiple substrates falling within the interaction distance in a single iteration. In these cases, a decision is made on which substrate to bind based on proximity. The risk of this can be minimised by reducing the time step accounting for the *k*_*f*_ and concentration of reactants. In this section we have shown new ABM methods and demonstrated that they are able to reproduce second-order reaction kinetics successfully across a wide range of scenarios.

### Reversible reactions

Reversible reactions form an integral part of many cell-signalling pathways, with dynamic forward and reverse reactions occurring simultaneously even in steady-state conditions. The combination of the methods described above for the different forward and reverse reactions can be combined together to give reversible reactions. However, when using a second order forward reaction, as reactions occur according to proximity, this can lead to a problem where two molecules are highly to react immediately after their dissociation, a phenomenon termed as germinate recombination in the work of Andrews and Bray [18]. We limit this effect by the introduction of an unbinding distance *d*_*u*_ equal to:

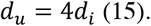

The unbinding distance is an arbitrary distance used to separate two reactive molecules after dissociation, thus preventing their immediate reassociation. Therefore, by combining this with the second order reaction and first order reaction methodology presented previously, we can model reversible reactions by ABM.

### Integration into the FLAME-accelerated Signalling Tool

Cell signalling networks involve complex networks of many reactants and reactions occurring simultaneously. Therefore, once we established and validated the methods described in the previous section we set out to create a tool for the facile writing of complex cell signalling networks as ABM simulations compatible with FLAME and FLAME GPU. The FaST is a Matlab-encoded tool fronted with a graphical user interface. It requires two text input files, one containing agent properties and the other containing reaction properties. The agent input file lists the agent name, its cellular localisation, diffusion coefficient and concentration. The reaction input file lists the reactants and products involved in each reaction, the type of reaction and the reaction constants. The FLAME-accelerated Signalling Tool produces FLAME and FLAME GPU simulation code from these input files along with associated tools for data retrieval and initial state generation. Furthermore, the FaST has the option to produce an equivalent ODE model for direct comparison between ABM and ODE simulations. The codes can be compiled as they are for CPUs or GPUs. However, they can also be easily modified to create new bespoke ABM codes for parallelisation on CPUs or GPUs, taking advantage of the inherent parallelisation optimisation in FLAME and FLAME GPU (Fig 2). Example input files are provided in the supplementary information with this paper and the tool itself is provided through the Zenodo platform (10.5281/zenodo.2620048).

**Fig 2.**
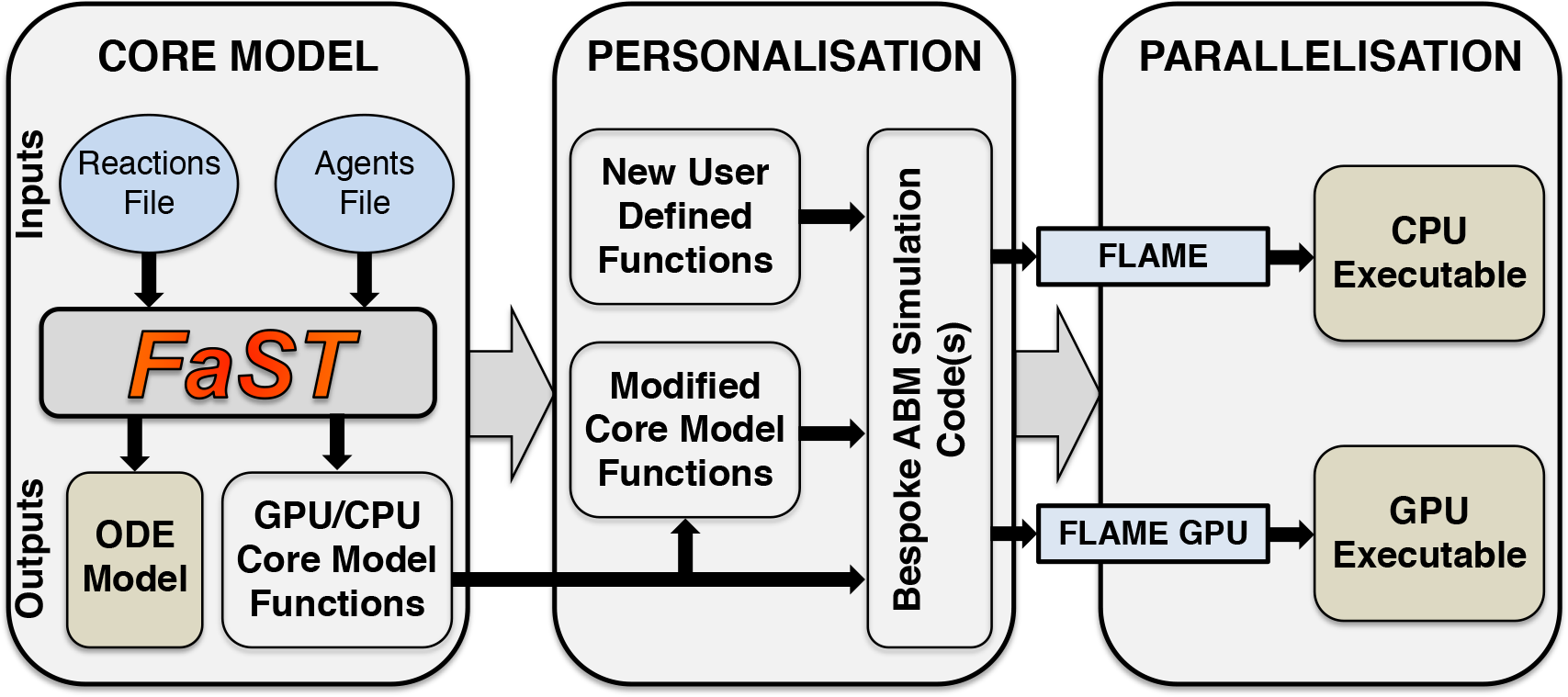
Schematic of the FLAME-accelerated Signalling Tool. The *Agents Input File* includes information on the localisation, concentration and diffusion coefficient of the agents in the system. The *Reactions Input File* includes the reaction, reaction type and reaction kinetic data. The FaST then generates ABM files compatible with FLAME and FLAME GPU, as well as an ODE model. FLAME and FLAME GPU then convert these into ABM executable models using the appropriate compilation tools. Furthermore, the user can also modify the core ABM code and add new functions, but still easily parallelise their bespoke ABM simulation code using FLAME or FLAME GPU.

### The FLAME-accelerated Signalling Tool is able to convert ODE models into agent-based models

In order to test the applicability of the FaST to modelling cell-signalling processes, we used it to convert a modified form of a previously well-characterised ODE model of apoptosis execution signalling into an ABM model [7]. In apoptosis execution signalling, upstream death signals lead to mitochondrial outer membrane permeabilisation (MOMP). This permeabilisation allows the release of the mitochondrial located proteins cytochrome c and Second Mitochondria-derived Activator of Caspases (SMAC). These two proteins activate a signalling cascade that results in apoptosis. This signalling requires the formation of a protein complex called the apoptosome [32,33]. This complex is used as a platform for the activation of the inactive precursor of the initiator caspase, pro-caspase 9 (PC9), into caspase 9 (C9), which in turn activates the inactive precursor of the executioner caspase, pro-caspase 3 (PC3), into the active caspase 3 (C3). C3 then cleaves numerous downstream substrates that invoke apoptosis, but can also cleave C9 [32–35]. This process is inhibited by the actions of X-linked Inhibitor of Apoptosis Protein (XIAP), which binds, inhibits and promotes ubiquitin-mediated degradation of the active form of C3 and C9 [36–39]. However, XIAP is unable to bind and inhibit C3-cleaved C9 (C9P) creating a positive-feed back loop. The actions of XIAP are countered by SMAC, which, after its release from the mitochondria, binds XIAP and actively breaks up caspase-XIAP complexes [40–43]. The reaction network consists of 14 protein/protein complexes and 23 individual reactions (Fig 3a). Full details of the model, including starting concentrations, reactions, reaction kinetics and diffusion coefficients are summarised in the supplementary material (supplementary S5). The model was placed into the setup text files required by the FaST, which are also included in the supplementary material (supplementary S6-7 Texts). The FaST was then used to make the ABM simulations for both FLAME and FLAME GPU.

**Fig 3.**
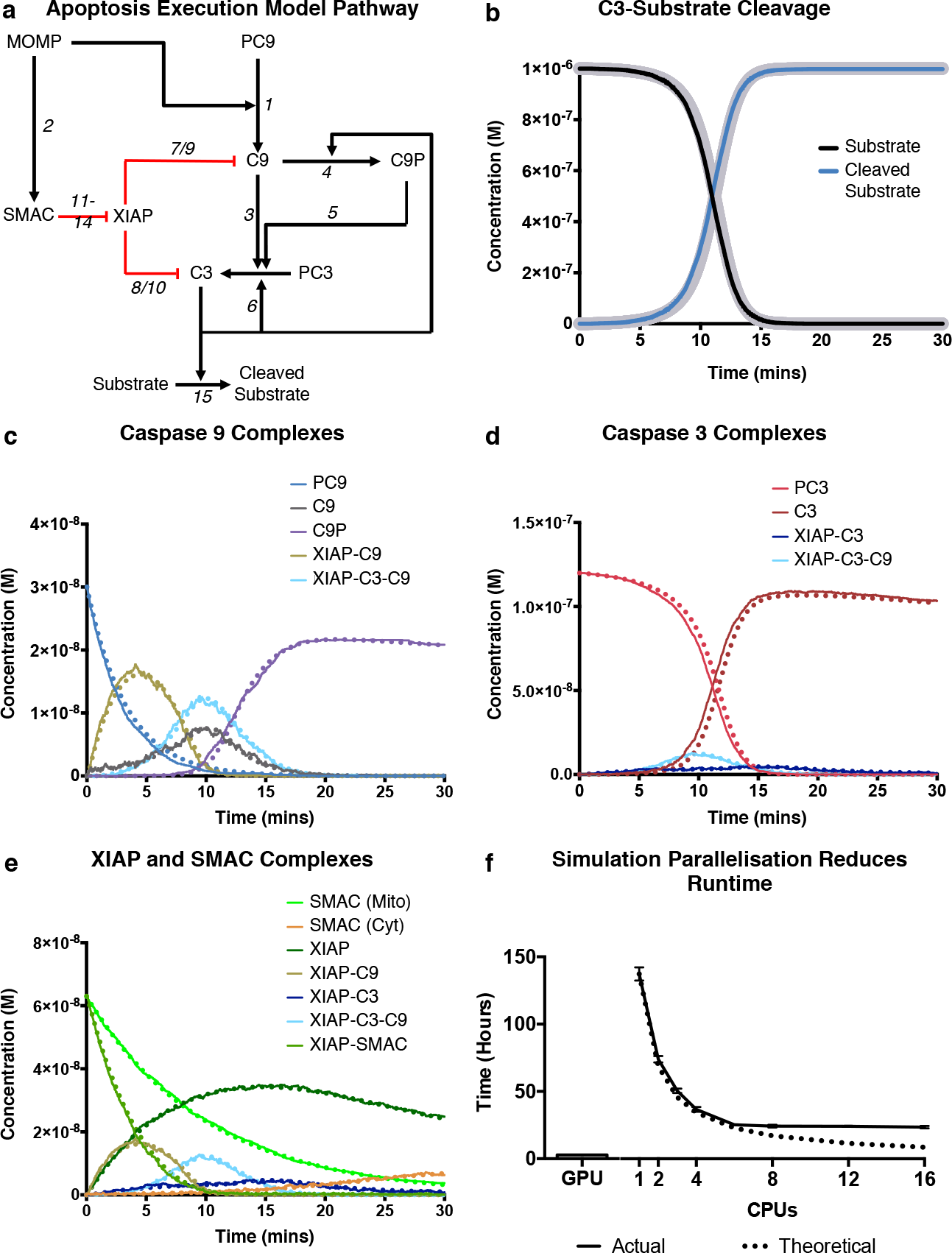
Agent-based models can be used to model complex signalling networks. The network of reactions in the model of apoptosis execution is shown (**a**). ABM simulations (thin solid lines) of apoptosis execution were compared to equivalent ODE simulations (thick grey lines for b; dashed lines for c-e) for final substrate cleavage (**b**); pro, activated or complex forms of caspase 9 (**c**); pro, activated or complex forms of caspase 3 (**d**); and XIAP or SMAC complexes (**e**). Run times of the simulation in **b-e** by FLAME GPU (bar) and by FLAME with increasing numbers of CPUs (solid line) is compared to the theoretical maximal speed up (dashed line) (**f**). All simulations were for 30 minutes, the time step *Δt* for particle diffusion was 0.0001 s and for reactions was 0.05 s in all simulations. Points in **f** represent mean ± standard deviation for three independent simulations.

The ABM simulations were run with >20000 agents and the progression of the reactions were compared, over 30 minutes, against the ODE model (Figs 3b-e). The apoptosis execution model culminates with the cleavage of a substrate of C3, to reflect the C3-mediated cleavage of an experimental Förster Resonance Energy Transfer (FRET) probe. The cleavage of this substrate showed excellent agreement between the ODE and ABM simulations with an R^2^ > 0.999 (Fig 3b). The activity of C9, C3 and the XIAP-SMAC regulatory axis in the ODE and ABM simulations is compared in Fig 3c, 3d and 3e respectively. All complexes within the ABM simulation show excellent agreement with the ODE simulation, demonstrating that the ABM methods used in the FaST can indeed reproduce mass action kinetics of ODE simulations.

### The FLAME-accelerated Signalling Tool can improve ABM performance by parallelisation

As the relatively poor performance of ABM simulations is the major drawback of ABM compared to other methods such as ODE, SDE or PDE models, we next looked at whether the use of GPU and CPU parallelisation could improve the speed of ABM simulations. We took the GPU and CPU versions of the apoptosis execution model and ran the simulation under the same conditions but on a GPU or on 1, 4, 6, 8, 12 or 16 CPUs in parallel, connected by 4× FDR InfiniBand interconnect. The total run time, in hours, for each simulation is displayed in Fig 3f and compared to the perfect theoretical speed up. The parallelisation of the ABM simulation on both GPUs and CPUs improved the runtime of the simulations compared to running on a CPU in serial with a speed up of 53.8× for GPU-acceleration and 8.9× for parallelisation across 16-CPUs. The speed up efficiency of CPU parallelisation can be calculated from the observed and theoretical speed up, for our ABM simulation this ranged from 55% (16-cores) to 62% (6-cores). Parallelisation efficiency decreased as more CPUs were added but this change was gradual suggesting that the addition of further CPUs would further increase speed up. Most notably, however, the GPU-accelerated version reduced the run time to under 3 hours.

## Discussion

ABM is a powerful method for modelling cellular signalling as it can include stochastic effects, heterogeneity and spatiotemporal organisation. The major challenges in ABM of cellular signalling is in increasing the scale of simulations, decreasing the time taken for simulation and producing robust accurate modelling of biological systems. The FaST, described and tested in this paper, is software that is able to produce ABM codes that robustly model biological systems and are parallelisable on GPUs or CPUs, thus addressing these limitations in ABM.

We extended previously established methods for ABM of cell signalling and validated their potential to reproduce mass action kinetics under a wide range of conditions [22]. These methods, based on the previous work of Pogson and colleagues, are similar in concept to those used in formal agent-based modelling software, such as Smoldyn, Chemcell and MCell, but operate at considerably longer time steps, therefore decreasing the amount of overall computation. Previously, it has been reported that the methods of Pogson and colleagues may not be able to reproduce kinetics observed under diffusion-limited, crowded or compartmentalised microenvironments [22,23]. We did not observe this effect in this paper (Figs 1 and 3) or in further testing (not shown), this is possibly due to the diffusion coefficients used in this paper and/or the simple geometry of the testing environment. In circumstances where this does become problematic, the correction suggested in the work of Klann and colleagues using collision rates is easily implementable in the framework we have presented [23]. In our ABM method, the reaction is driven entirely by proximity, rather than by collisions and activation energy that occur at the nanoscale. Several papers have suggested adapting the methodology to more accurately reflect these events using an increased binding radius and a probability of reaction [22,23]. The probability of reaction can be related to the activation energy, calculable from the rate constant through the Arrhenius equation. This is easily included within the methods presented here. However, such an inclusion would increase the amount of computation due to the increased binding radius and in most circumstances, this is unlikely to significantly change outcomes in the simulation.

We integrated the validated ABM methodology into the FaST, software that can take standard input files and generate ABM simulations compatible with FLAME and FLAME GPU. FLAME and FLAME GPU offer alternative approaches to parallelisation, increasing the feasibility of large-scale ABM simulations of cell signalling, as demonstrated in Fig 3f. The CPU parallelised version of the apoptosis model increased speed up of simulation by 8.9-fold when we used 16 CPUs. The speed up by CPU parallelisation is highly dependent on the amount of inter-agent communications, as it requires messaging through the message-passing interface. Agent-based modelling of cell signalling is communication heavy, as each individual protein (potentially millions in a single cell) has to communicate its own location, traditionally making it poorly suited for CPU parallelisation. Here we reported a speed-up efficiency of ~55-60% for parallelisation, compared to optimal speed up. The lag, caused from invoking the message passing interface, is dependent on the interconnect between individual CPU units. In this paper, we used a high-performance system using a 4× Fourteen Data Rate (FDR) InfiniBand interconnect. However, recent developments of Enhanced Data Rate (EDR) and High Data Rate (HDR) systems offer improved performance in interconnect, potentially reducing overheads associated with CPU parallelisation of agent-based applications.

Message heavy applications are generally more suited to GPUs, architecture specifically designed for massively parallel processing. We reported here that our GPU-accelerated ABM simulation of apoptosis increased the speed up of simulations by 53.9-fold, compared to the serial CPU version. This presents a major improvement in feasibility of undertaking ABM simulations of cell signalling pathways. Previously, GPU-implementations of the formal-simulator Smoldyn were shown to speed up simulations by up to 130-fold, although the GPU-implementation has reduced functionality compared to Smoldyn itself [44]. In most circumstances, GPU architecture likely represents the optimal platform for ABM simulations of cell signalling. However, GPUs are limited by their fixed amount of memory, which under certain circumstances may limit the scale of simulation compared to CPU versions of FLAME, where memory is less prohibitive [24][24]. One such scenario would be ABM-ODE hybrid simulations, for example, in a simulation where multiple cells undergo their own individual ODE for intracellular signalling, but simultaneously undertake intercellular signalling through ABM methods. Here, the memory required to store reactant concentrations for each individual cell may become impractical for GPUs but is well suited for CPU parallelisation.

The FaST is not designed to compete directly with formal simulators. Smoldyn and other formal agent-based modelling software packages offer optimised, accurate, user-friendly agent-based modelling with a wide-range of options in terms of geometric conditions [19–21]. The FaST, instead, is aimed to produce agent-based modelling code for simulating cell signalling, where greater personalisation and flexibility in functionality is required. The advantage of using FLAME and FLAME GPU is the ease-of-access to parallelisation without the requirement for detailed knowledge in MPI or CUDA coding respectively. Moreover, both software packages offer a *plug-and-play* approach to agent-based modelling, where additional functionality can be added or removed through the use of self-contained functions, but with the inherent parallelisation optimisation used within FLAME. Therefore, code produced by the FLAME-accelerated Signalling Tool can be easily altered whilst retaining the ability to be easily parallelised. Previously, FLAME and FLAME GPU have been used to model signalling processes including the NFκB pathway, *Escherichia coli* oxygen sensing and the mitogen-activated protein kinase pathway [45–47]. However, these models have always been based on relatively simple reaction networks as implementation requires extensive coding. The FaST offers easy creation of bespoke ABM simulations, of more extensive reaction networks, for FLAME and/or FLAME GPU.

In conclusion, we have presented methodology and a new software tool, the FLAME-accelerated Signalling Tool, for the building of agent-based models of cellular signalling that are flexible and malleable but still can be easily parallelised on CPUs or GPUs using FLAME and FLAME GPU respectively.

## Methods

### Software

FLAME xparser, message libraries (libmboard) and visualiser were obtained from github (https://github.com/FLAME-HPC/) with further documentation provided at www.flame.ac.uk. The FLAME GPU Software Development Kit (SDK) was downloaded from github (https://github.com/FLAMEGPU/) with further documentation provided at www.flamegpu.com. FLAME models and analysis scripts were built and tested using GCC 4.2.1 packaged through Xcode 8.3.3 developer tools, in conjunction with OpenMPI 2.0.2 MPI libraries for parallel compilation. FLAME GPU applications were produced using the FLAME GPU SDK using NVIDIA CUDA 9. The FLAME-accelerated signalling tool was constructed using the Graphic User Interface tools in MATLAB 2016b, license number 886886. The ODE models produced by the FaST were run on MATLAB 2016b.

### Test models

The test models in Fig 1 and S2-4 were produced in FLAME and ran in serial on a 4 GHz iMac Intel Core i7. All testing in Fig 1, Fig 3, S2-4 and S6 were performed in a 3 μm × 3 μm × 3 μm square test environment, with the bottom edge treated as a planar membrane. Concentrations of all reactants, including membrane-bound molecules, were calculated relative to the fixed volume. All boundaries within the testing environment are treated as reflective boundaries. All simulations presented in this paper used a time step for particle diffusion of 0.0001 s and 0.05 s for reactions.

### Speed testing

The simulations in Fig 3b-c and speed testing in Fig 3d were performed either on the Baden-Württemberg Tier 3 High Performance Computing Uni Cluster (bwUniCluster) for CPU simulations or on an NVIDIA GeForce 630 2 GB graphics card for GPU simulations. Simulation code was compiled using standard GNU compilers with the parallel FLAME message board libraries (libmboard) and message passing interface libraries (OpenMPI). Parallel CPU simulations were run on Intel Zeon E5-2670 processors with a 4× FDR InfiniBand interconnect. Partitioning was performed using a round robin approach offered by the FLAME software. Alternatively, FLAME is able to undergo geometric partitioning, where partitioning agents on separate CPUs is performed according to position. The theoretical run time for a given number (*N*) of CPUs (*CPU*_*N*_) was calculated relative to the run time when the number of CPUs is equal to 1 (N=1):

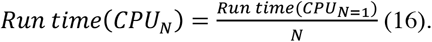

The speed up of parallelised simulations was calculated relative to the CPU serial model (*CPU*_*N=1*_) by the relation:

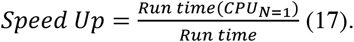

The efficiency of parallelisation on CPUs was calculated from the observed speed up and the theoretical speed up of *N*:

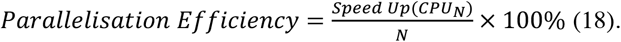

## Acknowledgements

The authors would like to thank Paul Richmond, Mozhgan Kabiri Chimeh and Peter Heywood at the University of Sheffield for support and feedback with using FLAME GPU.

## Availability

The FaST software tool including binaries, source code, template files and instructions for usage are available using the DOI: 10.5281/zenodo.2620048

