## supplementary S for "The FLAME-accelerated Signalling Tool (FaST): A tool for facile parallelisation of flexible agent-based models of cell signalling"

### Supplementary information for:
### Development of the FLAME-accelerated Signalling Tool: A tool for facile scale-up of flexible agent-based models of cell signalling

Gavin Fullstone*[,1,2], Cristiano Guttà[1], Amatus Beyer[1] and Markus Rehm*[,1,2]

#### S1 Text. FLAME: An approach to parallelisation of agent-based applications

In order to improve the scaling up of agent-based modelling, a number of different computational approaches are available. The scale of an agent-based simulation run on a CPU is limited by the amount of memory (primarily limits the number of agents) and computational power available (limits the number of agents and the functions that each agent can perform). In order to increase simulation size, distributed or parallel computing can be used, where memory and computation is spread across multiple processing units. This can be done with CPUs or alternatively with GPUs, such as those in computer graphics cards. GPUs usually offer greater computational power than CPUs and therefore are an attractive alternative for large-scale data processing, however they are limited by the fixed amount of memory. In agent-based modelling, this makes them very attractive for smaller, less memory intensive simulations but may limit their use in larger simulations or memory-intensive simulations such as ABM-ODE hybrids. Parallel computing with CPUs removes the theoretical upper limit of simulation size, based on memory and computational demands. CPU parallelisation uses the Message Passing Interface (MPI), which allows CPUs to send data to and receive data from other CPUs. However, whilst reading and writing locally stored data is relatively fast, messages passed through the MPI are comparatively slow and limited by the speed of the interconnect between CPUs. Therefore, careful optimisation of parallel codes is required to obtain maximum speed up of the simulation.

##### Agent-based modelling with FLAME

The major drawback of GPU and parallel CPU agent-based simulation is that it normally requires extensive knowledge of CUDA or MPI coding respectively and careful optimisation. FLAME and FLAME GPU use a communicating X-machine approach to agent-based modelling. The user defines agents; their respective memory variables; their respective functions to carry out and input/output requirements for those functions in the form of messages. This is achieved in a basic form using XMML (X-Machine Markup Language). The functions themselves are then coded in a separate function file or files coded in the C language. FLAME and the FLAME GPU Software Development Kit (SDK) utilises these two user-generated codes to construct iterative-based executable models in either for CPUs or GPUs respectively. FLAME and FLAME GPU use messages for communications between agents. The XMML file declares these messages and the variables stored in them. The CPU version of FLAME stores these messages on message boards. Functions that require data from other agents read through all the messages on a particular message board and by using filtering can access the required data efficiently. In serial, agents can read and write to this board rapidly as it is maintained in the local memory. However, in parallel each core has its own message board that must be synced with the other boards. When a function adds data to a message board, a message is generated to update the other message boards. Whilst messages are in transit through the message-passing interface, no function can read data from the message board. FLAME's approach to parallelisation uses a scheduler that prioritises work that will generate communications between CPUs. Whilst the messages are sent in the background, it will then perform all possible work not dependent on those communications, in order to reduce overhead times associated with parallelisation. This method of parallelisation optimisation has the additional advantage that the addition or removal of functions does not require re-optimisation and therefore allows a *plug and play* approach to adding functionality.

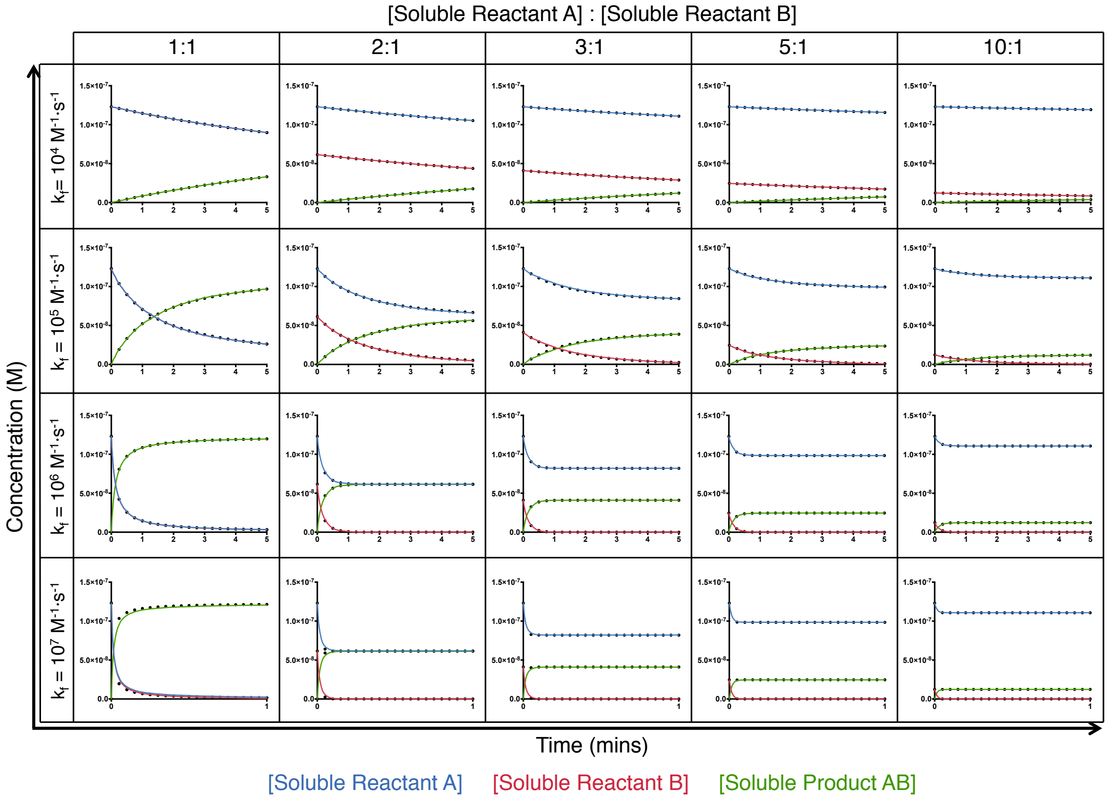


**S2 Fig. Agent-based modelling of soluble-soluble reactions is able to reproduce mass action kinetics.** The reaction of two soluble molecules were simulated with ABM and the change in reactant and product concentration was observed (solid colour lines) and compared to equivalent ODE models (dashed black lines) for $k_f$ values of $10^4$ $M^{-1}{\cdot}s^{-1}$, $10^5$ $M^{-1}{\cdot}s^{-1}$, $10^6$ $M^{-1}{\cdot}s^{-1}$ and $10^7$ $M^{-1}{\cdot}s^{-1}$ and different concentration ratios of A to B ([A]:[B]). All simulations were for 5 minutes, the time step $\Delta t$ for particle diffusion in all simulations was 0.0001 s and for reactions was 0.05 s. The diffusion coefficients used were 30 $\mu m^2{\cdot}s^{-1}$. Each point represents mean from three independent simulations, consequent error bars are too small to be plotted.

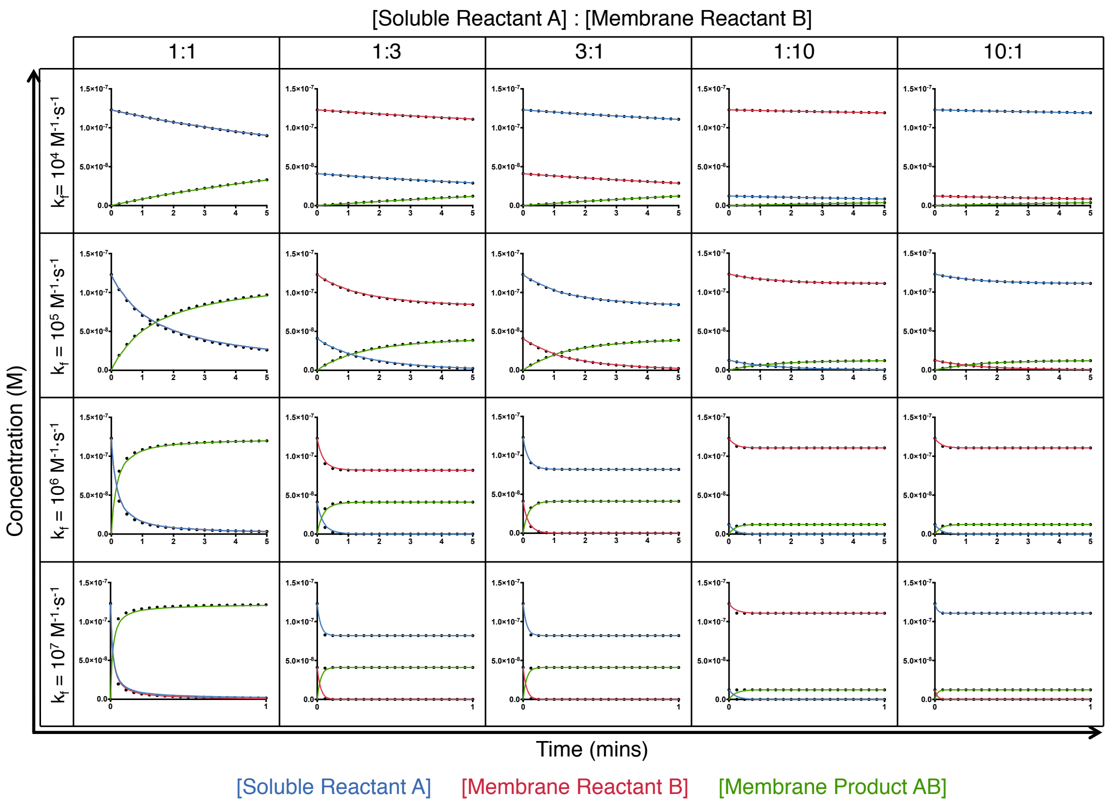


**S3 Fig. Agent-based modelling of membrane-soluble reactions is able to reproduce mass action kinetics.** The reaction of a soluble and a membrane-bound molecule were simulated with ABM and the change in reactant and product concentration was observed (solid colour lines) and compared to equivalent ODE models (dashed black lines) for $k_f$ values of $10^4$ $M^{-1}\cdot s^{-1}$, $10^5$ $M^{-1}\cdot s^{-1}$, $10^6$ $M^{-1}\cdot s^{-1}$ and $10^7$ $M^{-1}\cdot s^{-1}$ and different concentration ratios of A to B ([A]:[B]). All simulations were for 5 minutes, the time step $\Delta t$ for particle diffusion in all simulations was 0.0001 s and for reactions was 0.05 s. The diffusion coefficients used were 30 $\mu m^2\cdot s^{-1}$ for the soluble A and 0.3 $\mu m^2\cdot s^{-1}$ for the membrane-bound B. Each point represents the mean from three independent simulations, consequent error bars are too small to be plotted.

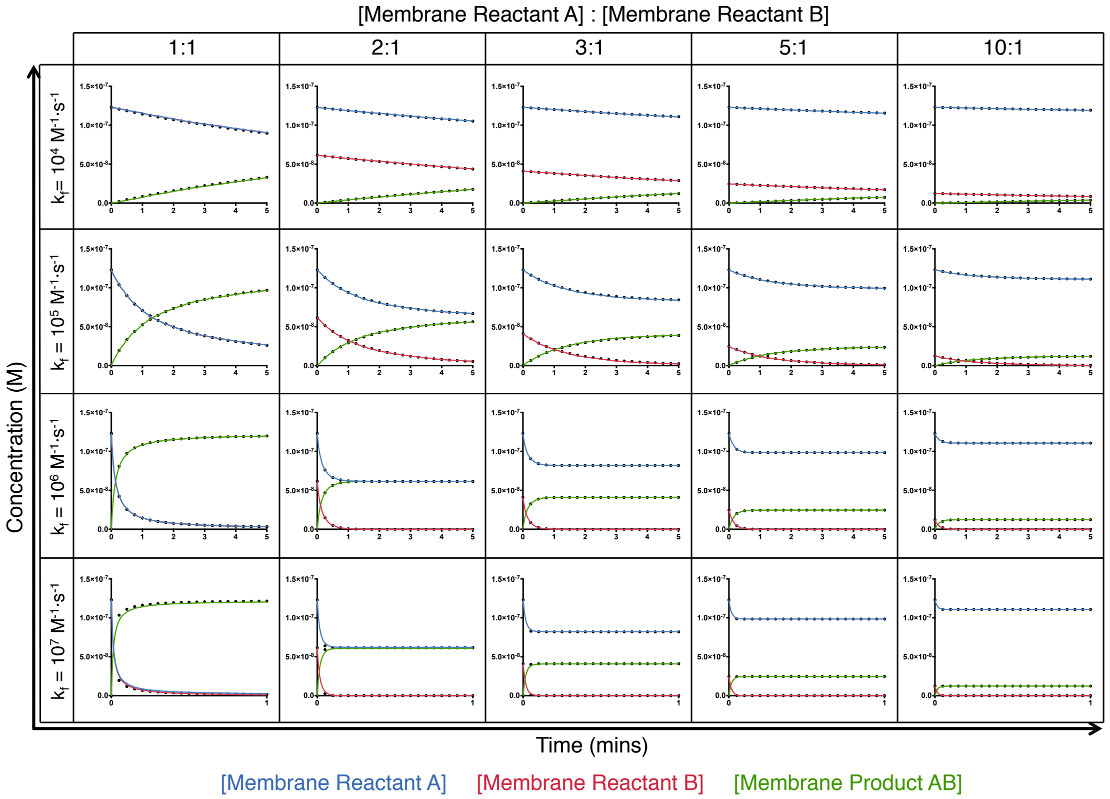


**S4 Fig. Agent-based modelling of membrane-membrane reactions is able to reproduce mass action kinetics.** The reaction of two membrane-bound molecules were simulated with ABM and the change in reactant and product concentration was observed (solid colour lines) and compared to equivalent ODE models (dashed black lines) for $k_f$ values of $10^4\ M^{-1} \cdot s^{-1}$, $10^5\ M^{-1} \cdot s^{-1}$, $10^6\ M^{-1} \cdot s^{-1}$ and $10^7\ M^{-1} \cdot s^{-1}$ and different concentration ratios of A to B ([A]:[B]). All simulations were for 5 minutes, the time step $\Delta t$ for particle diffusion in all simulations was 0.0001 s and for reactions was 0.05 s. The diffusion coefficients used were 0.3 $\mu m^2 \cdot s^{-1}$. Each point represents mean from three independent simulations, consequent error bars are too small to be plotted.

#### S5 Text. Reproduction of an established ODE model using ABM

The ODE model of apoptosis execution is a simplified representation of this pathway, where pro-caspase 9 is activated according to kinetic data representing an entire process involving cytochrome c release, resultant formation of a platform called the apoptosome and subsequent apoptosome-dependent pro-caspase 9 activation to caspase 9. We adapted this original model with two minor alterations to create the modified ODE and ABM models. First, SMAC was always treated as a dimer in its native form; therefore its concentration and the reactions were adjusted accordingly. Second, the cleavage of XIAP by caspase 3 was removed from the existing model. The resultant reaction schematic modelled in this paper is included in Fig 3a. We took the original concentrations and reaction kinetic data from the work of Rehm and colleagues, as summarised in S5 Tables 1-2, to construct the agents.txt and reactions.txt files required by the FLAME signalling tool. These files are also included within the downloadable supplementary information.

**S5 Table 1.** List of Agents and Concentrations

| ID | Name | Concentration (M) | Diffusion Coefficient ($m^2 \cdot s^{-1}$) | Description |
|---|---|---|---|---|
| 1 | PC9 | $3\times10^{-8}$ | $2.024\times10^{-11}$ | Pro-Caspase 9 |
| 2 | C9 | 0 | $2.024\times10^{-11}$ | Caspase 9 |
| 3 | C9P | 0 | $2.462\times10^{-11}$ | Processed Caspase 9 |
| 4 | PC3 | $1.2\times10^{-7}$ | $2.201\times10^{-11}$ | Pro-Caspase 3 |
| 5 | C3 | 0 | $1.860\times10^{-11}$ | Caspase 3 |
| 6 | XIAP | $6.3\times10^{-8}$ | $1.893\times10^{-11}$ | XIAP |
| 7 | MitoSMAC | $6.3\times10^{-8}$ | $2.462\times10^{-11}$ | Mitochondrial SMAC |
| 8 | SMAC | 0 | $2.461\times10^{-11}$ | Cytoplasmic SMAC |
| 9 | XIAP-C3 | 0 | $1.490\times10^{-11}$ | XIAP-Caspase 3 |
| 10 | XIAP-C9 | 0 | $1.551\times10^{-11}$ | XIAP-Caspase 9 |
| 11 | XIAP-C3-C9 | 0 | $1.331\times10^{-11}$ | XIAP-Caspase 3-Caspase 9 |
| 12 | XIAP-SMAC | 0 | $1.671\times10^{-11}$ | XIAP-SMAC |
| 13 | Substrate | $1.00\times10^{-6}$ | $1.905\times10^{-11}$ | C3 Substrate |
| 14 | cSubstrate | 0 | $2.400\times10^{-11}$ | C3-Cleaved Substrate |

All concentrations were taken from the publication of Rehm et al., 2006. SMAC was here considered constitutively a dimer and so the concentration was halved accordingly [1]. Diffusion coefficients were calculated relative to GFP (27 kDa; $2.4\times10^{-11}$ $m^2 \cdot s^{-1}$) according to the methods described.

**S5 Table 2.** List of Reactions and Rates

| # | Reaction | Forward Rate | Reverse Rate |
|---|---|---|---|
| 1 | PC9 → C9 | $5.022\times10^{-3}$ $s^{-1}$ | |
| 2 | MitoSMAC → SMAC | $1.65\times10^{-3}$ $s^{-1}$ | |
| 3 | C9 + PC3 → C9 + C3 | $1\times10^{5}$ $M^{-1}\cdot s^{-1}$ | |
| 4 | C9 + C3 → C9P + C3 | $2\times10^{5}$ $M^{-1}\cdot s^{-1}$ | |
| 5 | C9P + PC3 → C9P + C3 | $8\times10^{5}$ $M^{-1}\cdot s^{-1}$ | |
| 6 | C3 + PC3 → C3 + C3 | $4\times10^{4}$ $M^{-1}\cdot s^{-1}$ | |
| 7 | C9 + XIAP ↔ XIAPC9 | $2.6\times10^{6}$ $M^{-1}\cdot s^{-1}$ | $2.4\times10^{-3}$ $s^{-1}$ |
| 8 | C3 + XIAP ↔ XIAPC3 | $2.6\times10^{6}$ $M^{-1}\cdot s^{-1}$ | $2.4\times10^{-3}$ $s^{-1}$ |
| 9 | C9 + XIAPC3 ↔ XIAPC3C9 | $2.6\times10^{6}$ $M^{-1}\cdot s^{-1}$ | $2.4\times10^{-3}$ $s^{-1}$ |
| 10 | C3 + XIAPC9 ↔ XIAPC3C9 | $2.6\times10^{6}$ $M^{-1}\cdot s^{-1}$ | $2.4\times10^{-3}$ $s^{-1}$ |
| 11 | XIAP + SMAC ↔ XIAPSMAC | $7\times10^{6}$ $M^{-1}\cdot s^{-1}$ | $2.21\times10^{-3}$ $s^{-1}$ |
| 12 | XIAPC9 + SMAC → XIAPSMAC + C9 | $7\times10^{6}$ $M^{-1}\cdot s^{-1}$ | |
| 13 | XIAPC3 + SMAC → XIAPSMAC + C3 | $7\times10^{6}$ $M^{-1}\cdot s^{-1}$ | |
| 14 | XIAPC3C9 + SMAC → XIAPSMAC + C3 | $7\times10^{6}$ $M^{-1}\cdot s^{-1}$ | |
| 15 | C3 + Substrate → C3 + cSubstrate | $2\times10^{5}$ $M^{-1}\cdot s^{-1}$ | |
| 16 | C9 → DEGRADED | $9.67\times10^{-5}$ $s^{-1}$ | |
| 17 | C3 → DEGRADED | $9.67\times10^{-5}$ $s^{-1}$ | |
| 18 | C9P → DEGRADED | $9.67\times10^{-5}$ $s^{-1}$ | |
| 19 | XIAPC9 → DEGRADED | $5.78\times10^{-4}$ $s^{-1}$ | |
| 20 | XIAPC3 → DEGRADED | $5.78\times10^{-4}$ $s^{-1}$ | |
| 21 | XIAPC3C9 → DEGRADED | $5.78\times10^{-4}$ $s^{-1}$ | |
| 22 | XIAPSMAC → DEGRADED | $5.78\times10^{-4}$ $s^{-1}$ | |
| 23 | SMAC → DEGRADED | $9.67\times10^{-5}$ $s^{-1}$ | |

All reaction rates were used as in the publication of Rehm *et al.*, 2006, except for the reaction rates 11-14 which were taken from Huang *et al.*, 2003 [14]. The reaction rate for equations 1-2 were converted from half times in Rehm *et al.*, 2006 by the relation $k = 0.693/t_{1/2}$ where $t_{1/2}$ is the half time [1].
